# Sequence evaluation and comparative analysis of novel assays for intact proviral HIV-1 DNA

**DOI:** 10.1101/2020.10.07.330787

**Authors:** Christian Gaebler, Shane D. Falcinelli, Elina Stoffel, Jenna Read, Ross Murtagh, Thiago Y. Oliveira, Victor Ramos, Julio C. C. Lorenzi, Jennifer Kirchherr, Katherine S. James, Brigitte Allard, Caroline Baker, JoAnn D. Kuruc, Marina Caskey, Nancie M. Archin, Robert F. Siliciano, David M. Margolis, Michel C. Nussenzweig

## Abstract

The HIV proviral reservoir is the major barrier to cure. The predominantly replication-defective proviral landscape makes the measurement of virus that is likely to cause rebound upon ART-cessation challenging. To address this issue, novel assays to measure intact HIV proviruses have been developed. The Intact Proviral DNA Assay (IPDA) is a high-throughput assay that uses two probes to exclude the majority of defective proviruses and determine the frequency of intact proviruses, albeit without sequence confirmation. Quadruplex PCR with four probes (Q4PCR), is a lower-throughput assay that uses limiting dilution long distance PCR amplification followed by qPCR and near-full length genome sequencing (nFGS) to estimate the frequency of sequence-confirmed intact proviruses and provide insight into their clonal composition. To explore the advantages and limitations of these assays, we compared IPDA and Q4PCR measurements from 39 ART-suppressed people living with HIV. We found that IPDA and Q4PCR measurements correlated with one another but frequencies of intact proviral DNA differed by approximately 19-fold. This difference may be in part due to inefficiencies in long distance PCR amplification of proviruses in Q4PCR, leading to underestimates of intact proviral frequencies. In addition, nFGS analysis within Q4PCR explained that some of this difference is explained by proviruses that are classified as intact by IPDA but carry defects elsewhere in the genome. Taken together, this head-to-head comparison of novel intact proviral DNA assays provides important context for their interpretation in studies to deplete the HIV reservoir and shows that together the assays bracket true reservoir size.

**Importance:** The Intact Proviral DNA Assay (IPDA) and Quadruplex PCR (Q4PCR) represent major advances in accurately quantifying and characterizing the replication competent HIV reservoir. This study compares the two novel approaches for measuring intact HIV proviral DNA in samples from 39 ART-suppressed people living with HIV, thereby informing ongoing efforts to deplete the HIV reservoir in cure-related trials.

## Introduction

Latent replication-competent HIV proviruses are a formidable barrier to HIV cure, but there is little consensus on how to best measure this reservoir (1). Available assays for the reservoir include measurements of HIV DNA, inducible HIV RNA/protein, and quantitative viral outgrowth assays (2). Comparisons of different reservoir measurements provide useful insights into the nature of the HIV reservoir. For example, comparison of reservoir measurements revealed a greater than two log discrepancy between the frequency of integrated proviruses measured by DNA PCR and replication-competent proviruses measured by viral outgrowth (3). This discrepancy is largely explained by the disproportionate frequency of defective proviral DNA and variable inducible outgrowth of intact proviruses (4, 5). Thus, single-probe HIV DNA PCR based assays are limited in specificity for intact HIV genomes because a large fraction of the proviruses that they detect are defective.

Although quantitative viral outgrowth assays are specific for intact HIV genomes, they are labor intensive and the results vary over time by as much as a factor of 6 (6). Furthermore, they underestimate the frequency of intact HIV genomes because not all proviruses are equally inducible in vitro (5, 7). As such, many intact proviruses are not detected in outgrowth assays. To improve on the specificity of PCR-based assays for detection of intact proviruses novel methods have been developed that combine probes that target relatively conserved regions of the genome (Intact Proviral DNA Assay [IPDA]) (8) in addition to limiting dilution near-full genome proviral sequencing (a component of Q4PCR) (9).

The IPDA utilizes digital droplet PCR (ddPCR) to measure proviruses with probes targeting conserved regions in the packaging signal (PS) and envelope (*env*) that when amplified together in the same provirus, exclude the majority of defective proviruses. Input cells are directly measured by a simultaneous ddPCR reaction, enabling the IPDA to report the total frequency of intact and defective proviruses per million input cells. The Q4PCR assay employs long distance PCR at limiting dilution to amplify proviruses, followed by interrogation with four quantitative PCR (qPCR) probes for PS, *env*, *pol*, and *gag* in a multiplex reaction to detect amplified HIV-1 genomes. Cell inputs are estimated by quantification of input DNA and the frequency of intact proviruses are reported after sequence verification by near full-length genome sequencing (nFGS). Because the Q4PCR employs limiting dilution near full-length proviral amplification, these nFGS results can be used to provide insight into clonal composition of intact proviruses.

Here we compare quantitative viral outgrowth assays, Q4PCR, IPDA and total HIV gag DNA measurements on samples from 39 ART-suppressed people living with HIV (PLWH) to explore the advantages and limitations of these assays.

## Results

### Quantitative Comparison

Leukapheresis or whole blood samples were obtained from PLWH, stably suppressed (< 50 copies of HIV-1 RNA/ml) on ART, under protocols approved by the University of North Carolina (UNC) or Rockefeller University biomedical institutional review boards. The majority of study participants were treated during chronic infection and had been on ART for a median of 8.4 years (Table S1). For all 39 participants, Q4PCR and IPDA were performed on DNA extracted from total CD4 (tCD4) T cells from these two cohorts. Total HIV gag DNA was measured in tCD4 T cells using an assay designed to capture group M HIV-1 proviral DNA (10). Finally, we performed Q^2^VOA (n = 11, Rockefeller cohort, tCD4 cells) or QVOA (n = 13, UNC cohort, resting CD4 T cells) to measure inducible replication-competent HIV.

As expected, the sum of the IPDA-derived intact, 3’-defective, and 5’-defective proviral frequencies (IPDA Total) yielded the highest proviral frequency estimates (median 652 copies/million CD4 T cells), and quantitative viral outgrowth assays the lowest across both cohorts (median 0.60 infectious units/million CD4 T cells) (Fig. 1A). Median proviral frequency using a single probe LTR-*gag* assay (*gag* DNA) was 387 copies/million CD4 T cells. As expected, total HIV DNA frequency estimates (IPDA total DNA and *gag* DNA), were strongly correlated (Spearman r = 0.86, p = 0.001).

**Fig. 1:**
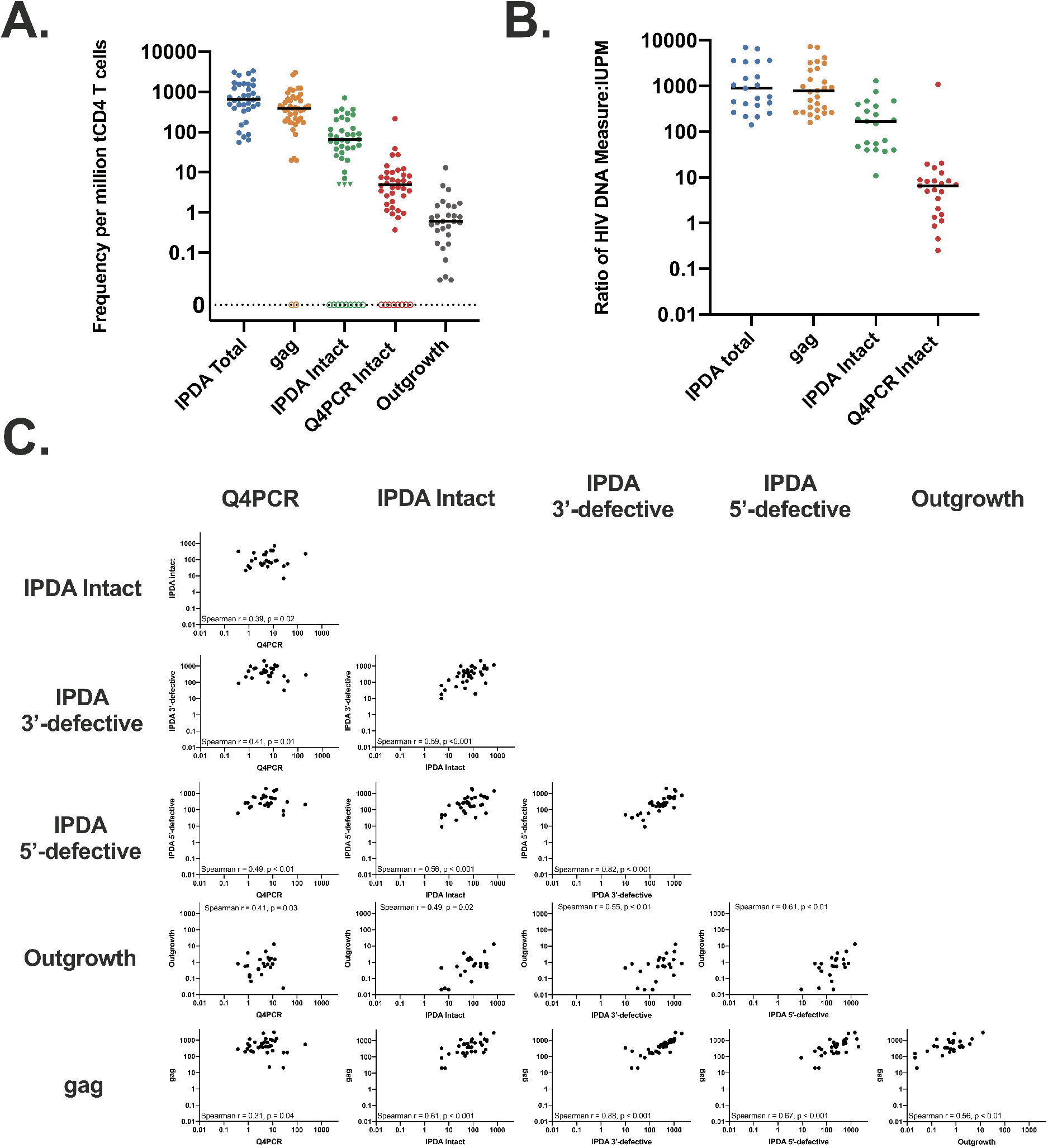
Quantitative Comparison of Reservoir Measurements. (A) Frequency per million total CD4 T cells for total HIV gag, intact provirus (IPDA), intact provirus (Q4PCR), and replication-competent outgrowth viruses. Triangles indicate left-censored measurements for the IPDA (detectable defective proviruses but no PS+*env+* proviruses, see Methods). Assay data from participants with an amplification failure (7/39 for IPDA, 2/39 for *gag*) or no recovery of intact proviral sequences for Q4PCR (8/39) are represented as hollow symbols and were excluded from the analysis, but other assay data for those participants was included if available. (B) Frequency ratio with viral outgrowth measures (IUPM) as the denominator for total HIV gag, intact provirus (IPDA), and intact provirus (Q4PCR). Data points that were left-censored, had an IPDA amplification failure, or had no recovered Q4PCR intact proviral sequences were excluded for this analysis. (C) Scatterplots showing correlations of reservoir measurements including Spearman r and unadjusted p-values. Left censored measurements were included: for Q4PCR, when no intact proviruses were recovered a value of 0 was used and for IPDA, a left-censor of 5 copies/million CD4 T cells was used as described in the Methods. Data points were excluded when an IPDA amplification failure was present.

The frequency of intact HIV genomes detected by IPDA and Q4PCR were intermediate between *gag* DNA and viral outgrowth, consistent with previous studies with these assays and with previous estimates based on limiting dilution proviral nFGS (4, 8, 9, 11). For IPDA, proviruses were considered intact if they were positive for both PS and *env* probes. For Q4PCR, proviruses were considered intact if they were sequence verified. The median intact proviral frequency per million CD4 T cells was 65 for IPDA and 5 for Q4PCR (Fig. 1A). IPDA measurements were a median of 19 (interquartile range 7, 48) fold higher than Q4PCR measurements. Compared to outgrowth assays, the median ratio of gag DNA (779), intact sequences by IPDA (169), or Q4PCR (7) was greater than one (Fig. 1B).

To further understand the relationship between the assays we calculated the Spearman correlation between gag, IPDA intact, IPDA 3’-defective, IPDA 5’-defective, IUPM, Q4PCR Intact (Fig. 1 C and Fig. S1). IPDA and Q4PCR measures of intact DNA showed similar correlations with quantitative viral outgrowth measures (Spearman r = 0.49 and 0.41, respectively), and with each other (Spearman r = 0.39). IPDA and Q4PCR both moderately correlated with gag DNA (r = 0.61 and r = 0.31, respectively), in agreement with a previous report comparing IPDA and *gag* measurements (12). 5’-defective and 3’-defective proviral frequencies measured using the IPDA also moderately correlated with quantitative viral outgrowth, IPDA intact, and Q4PCR proviral frequencies (Figure 1 C and Fig. S1).

### IPDA and Q4PCR Proviral Amplification

We observed amplification signal failure of either the PS or *env* IPDA primer/probe set for 7 (18%) of the 39 participants (0 of 12 from UNC cohort, and 7 of 27 from Rockefeller cohort). In addition, another small subset of samples (4 of 39 - 10.3%) showed reduced fluorescent signal intensities in the PS or *env* IPDA primer/probe set thereby complicating the unambiguous identification of a distinct double positive droplet population. With Q4PCR we were able to identify sequences that were positive for 2 or more probes in 37 of the 39 individuals (9). However, we did not detect intact sequences from 8 of the 39 participants (20.5%).

We determined viral genotypes using the Q4PCR sequencing data. Overall, we identified 3 participants (one from UNC cohort, two from Rockefeller cohort) with non-subtype B viral infection (Clade A1 and G). For 2 of the 3 non-subtype B infected individuals we observed IPDA amplification failures in the *env* IPDA probe, reflecting the fact that the initial version of the IPDA was designed based on clade B sequence data (8). In contrast, using Q4PCR we retrieved intact proviral sequences from 2 of the 3 individuals with non-subtype B viruses. The utilization of 4 instead of 2 probes appears to render Q4PCR less sensitive to subtype variability. For example, *env* amplification failed for participant 5112 in both assays, but several intact proviral genomes scored positive for the PS, pol and gag probe by Q4PCR (Fig. S2).

For the fraction of participants harboring clade-B viruses, the frequency of IPDA amplification signal failure was 14.7% which is similar to but slightly higher than that reported by others in larger cohorts (11) (Fig. 2A and Table S2).

**Fig. 2:**
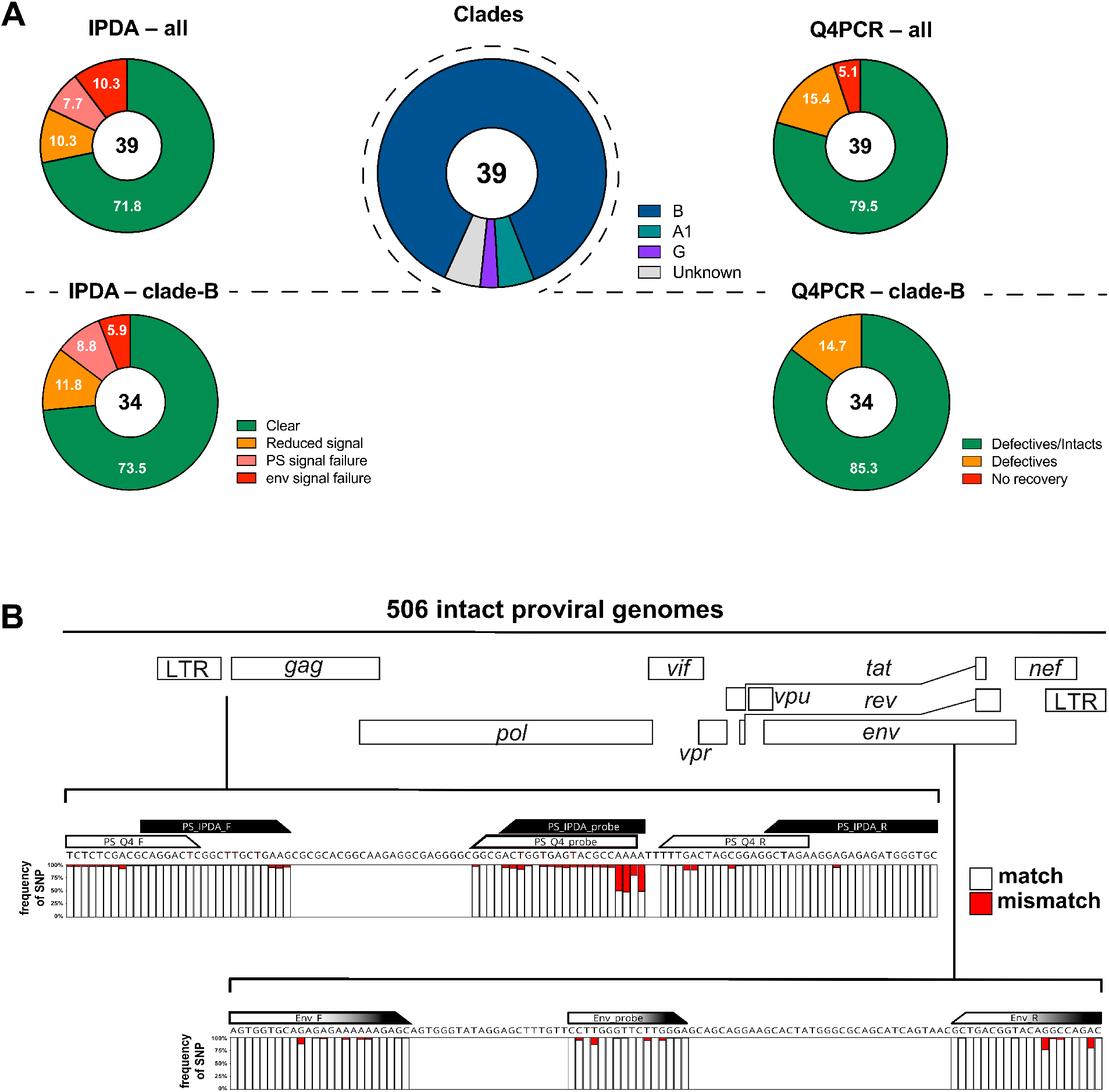
(A) IPDA and Q4PCR signal summary. Pie charts on the left depict the fraction of participants that showed an IPDA amplification failure for PS (light red slice) or *env* (dark red slice), reduced signal intensities/droplet separation (orange slice) or clear results (green slice) for all viral genotypes (upper panel) or only clade-B viruses (lower panel), respectively. Similarly, pie charts on the right depict the fraction of participants that yielded no sequence information (red slice), only defective proviral sequences (orange slice) or both defective and intact sequences (green slice) for all viral genotypes (upper panel) or only clade-B viruses (lower panel), respectively. Pie chart in the center shows the distribution of viral genotypes among the 39 participants (clade B (blue), A1 (green), G (purple) or unknown (grey). **(B) Sequence conservation in IPDA and Q4PCR PS+env primer/probe binding regions among 506 intact proviral genomes.**Stacked bar graphs show sequence identity obtained by aligning all intact proviral sequences with the PS primer/probe set (IPDA PS primer/probe set in black, Q4PCR PS primer/probe set in white, respectively) and *env* primer/probe set (identical for IPDA and Q4PCR, depicted in black and white). White or red bars represent the frequencies of single-nucleotide polymorphisms (SNPs) that match (white) or mismatch (red) the primer/probe in a specific nucleotide, respectively.

### PS and *env* sequence conservation among intact proviruses

Overall, we found 506 intact and 4211 defective proviral sequences by Q4PCR (Table S3). The IPDA and Q4PCR utilize overlapping primer/probe sets in PS and identical primer/probe in *env (9)*. To determine the level of sequence conservation in the primer/probe binding regions in our study participants at a single nucleotide resolution we aligned all 506 intact proviral genomes from Q4PCR nFGS.

As expected, the binding regions of the IPDA/Q4PCR PS and *env* primer/probe sets showed high levels of conservation among the intact proviral genomes (8, 9). Nevertheless, single nucleotide polymorphisms (SNP) could be detected to varying degrees in all primers and probes of both PS and *env*. While the occurrence of common SNPs in the 5’ region of primers/probes (e.g. up to 50% in the 5’ PS probe region) are expected to have little impact on signal quality, a mismatch in the very 3’ end of the primers/probes (e.g. 6% SNP frequency in the IPDA PS forward primer 3’ end) can impact signal detection considerably (13, 14) (Fig. 2B).

### PS and *env* sequence polymorphism analysis

To examine the importance of HIV diversity on an individual level, we next used the Q4PCR sequence information to study polymorphisms and associated signal patterns for individual participants.

In most cases the quality of IPDA and Q4PCR signal could be explained by absence or presence of sequence polymorphisms in the primer/probe binding regions of proviral genomes. For example, intact proviruses from participant UNC-434 showed complete conservation in the primer/probe binding regions resulting in clear and distinct IPDA and Q4PCR PS and *env* signals (Fig. 3A). In contrast, the amplification failure of the IPDA PS signal in participant 9243 could be explained by a two-nucleotide mismatch at the 3‘end of the PS IPDA forward primer (Fig. 3B). In addition, single nucleotide polymorphisms in the *env* primer/probe region of intact proviruses from participant 5101 (Fig. 3C) resulted in signal reduction rather than the complete amplification failure for the IPDA. While such a decrease in droplet separation complicates the clear identification of PS+*env* positive proviruses by IPDA, PCR annealing temperature decreases were shown to improve droplet resolution (Fig. S3). Proviral signal patterns (droplet amplitude or qPCR amplification curves) were found to be stable over time in all six individuals that were assayed at more than one time point, thereby demonstrating the consistent impact of sequence polymorphisms on signal patterns. However, occasional discordant results between qPCR amplification and sequencing results require further study.

**Fig. 3:**
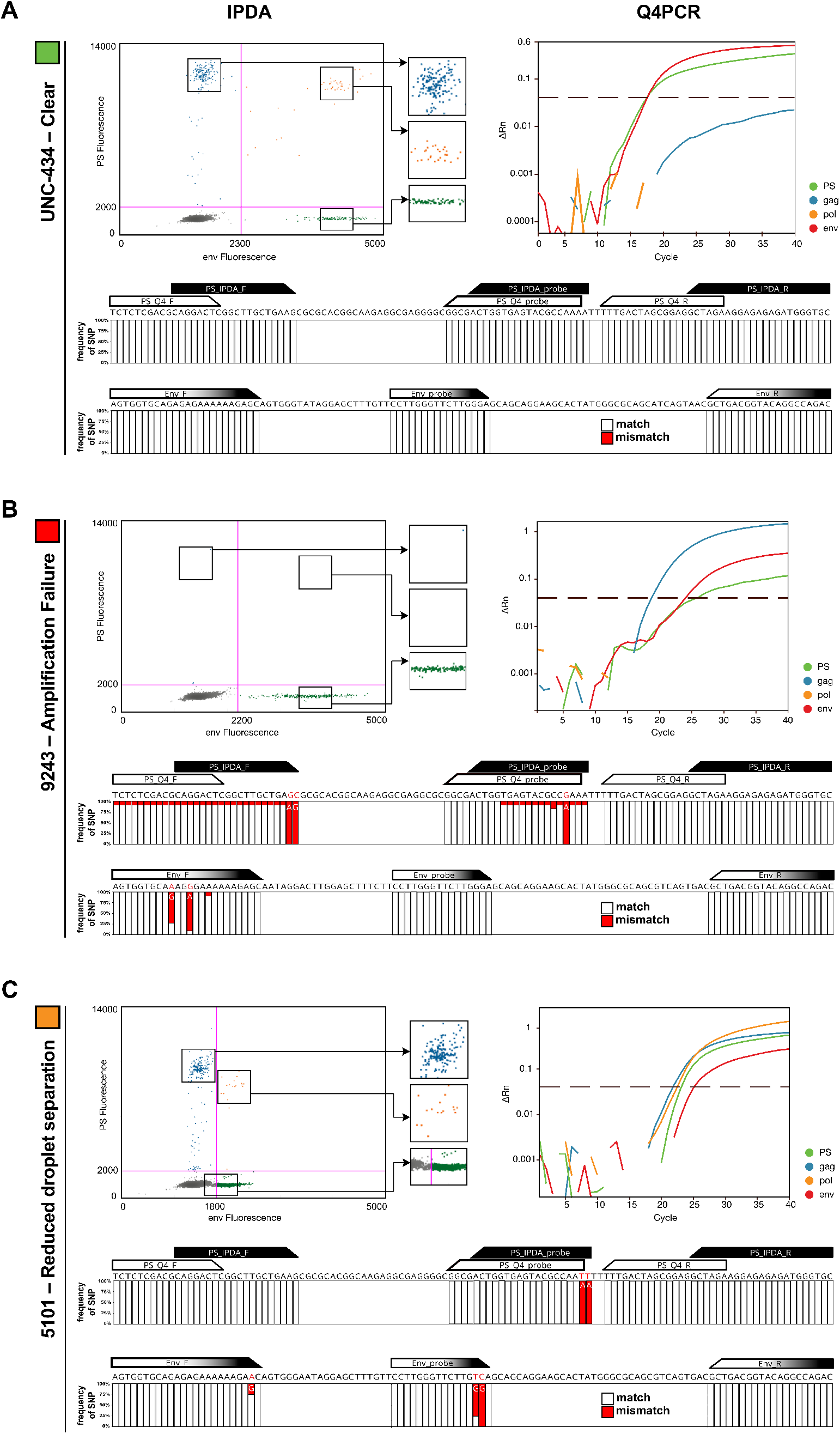
PS and env sequence polymorphism analysis. **(A)** Example of a clear droplet distribution for the PS (blue), *env* (green) and PS+*env*-positive (orange) droplets by IPDA and the corresponding Q4PCR signal (PS=green, *gag*=blue, *pol*=yellow, *env*=red) for participant UNC-434. Stacked bar graphs depict sequence alignment of all intact proviral sequences from participant UNC-434 with the PS primer/probe set (IPDA PS primer/probe set in black, Q4PCR PS primer/probe set in white, respectively) and *env* primer/probe set (identical for IPDA and Q4PCR, depicted in black and white). White or red bars represent the frequencies of single-nucleotide polymorphisms (SNPs) that match (white) or mismatch (red) the primer/probe in a specific nucleotide, respectively. **(B)** Example of an amplification failure of the PS signal by IPDA for participant 9243. Corresponding Q4PCR signal and intact proviral genome alignment reveals an explanatory two-nucleotide mismatch in the IPDA PS forward primer 3’ end across all intact proviral genomes from participant 9243. **(C)** Example of a reduced droplet distribution in the IPDA *env* signal thereby complicating the clear separation of the PS+*env*-positive droplets from the negative population for participant 5101. The corresponding Q4PCR signal shows a reduced signal intensity for the *env* signal as well. The intact proviral genome alignment depicts an explanatory two-nucleotide mismatch in the IPDA/Q4PCR probe 3’ end.

### PS and *env* intact proviral genome prediction

Intact proviral DNA detection assays face the challenges of both detecting viral sequences and correctly characterizing them as intact or defective. Using nFGS data, Bruner and colleagues reported that the IPDA PS and *env* primer/probe sets exclude 97% of proviruses with defects detectable by nFGS, and that approximately 70% of proviruses classified as intact by IPDA lack defects detectable by nFGS (8).

We used Q4PCR sequencing data to assess the fraction of true intact sequences out of all samples that were amplified by the Q4PCR PS and *env* primer/probe set combination. We considered a total of 4717 proviruses collected from all wells positive for 2 or more of the quadruplex qPCR reactions of the Q4PCR. Notably, proviral sequence was not collected from wells positive for only one qPCR reaction in the Q4PCR, which likely contain defective proviruses (9). Twenty-nine PS+*env*-positive sequences with a threshold cycle Ct>38 were excluded. Out of the remaining 4688 proviral genomes, only 625 or 13.3% scored positive with PS+*env* (Ct<38 for both amplicons). Out of 4139 defective proviruses positive for 2 or more of the quadruplex qPCR reactions in Q4PCR, only 305 (7.4%) were found to be positive for the combination of PS+*env* thereby demonstrating the exceptional specificity of this particular probe combination to exclude defective proviruses (9). At the same time, the Q4PCR PS+*env* combination failed to detect 175 (35.3%) out of 475 intact proviruses. The majority of the 175 PS+*env*-negative intact genomes were recovered from individuals with amplification signal failures of either the PS or *env* primer/probe set and only the utilization of two additional probes (*gag* and *pol*) enabled the detection and intact sequence verification by Q4PCR (Fig. 4A). Signal failure was generally consistent between Q4PCR and IPDA, except where there were signal-eliminating polymorphisms in the Q4PCR but not the IPDA PS primer/probe set or vice-versa (Fig. 3B). Importantly, amplicon signal failure in the IPDA is readily apparent, and intact provirus values are not reported in these cases (11).

**Fig.4:**
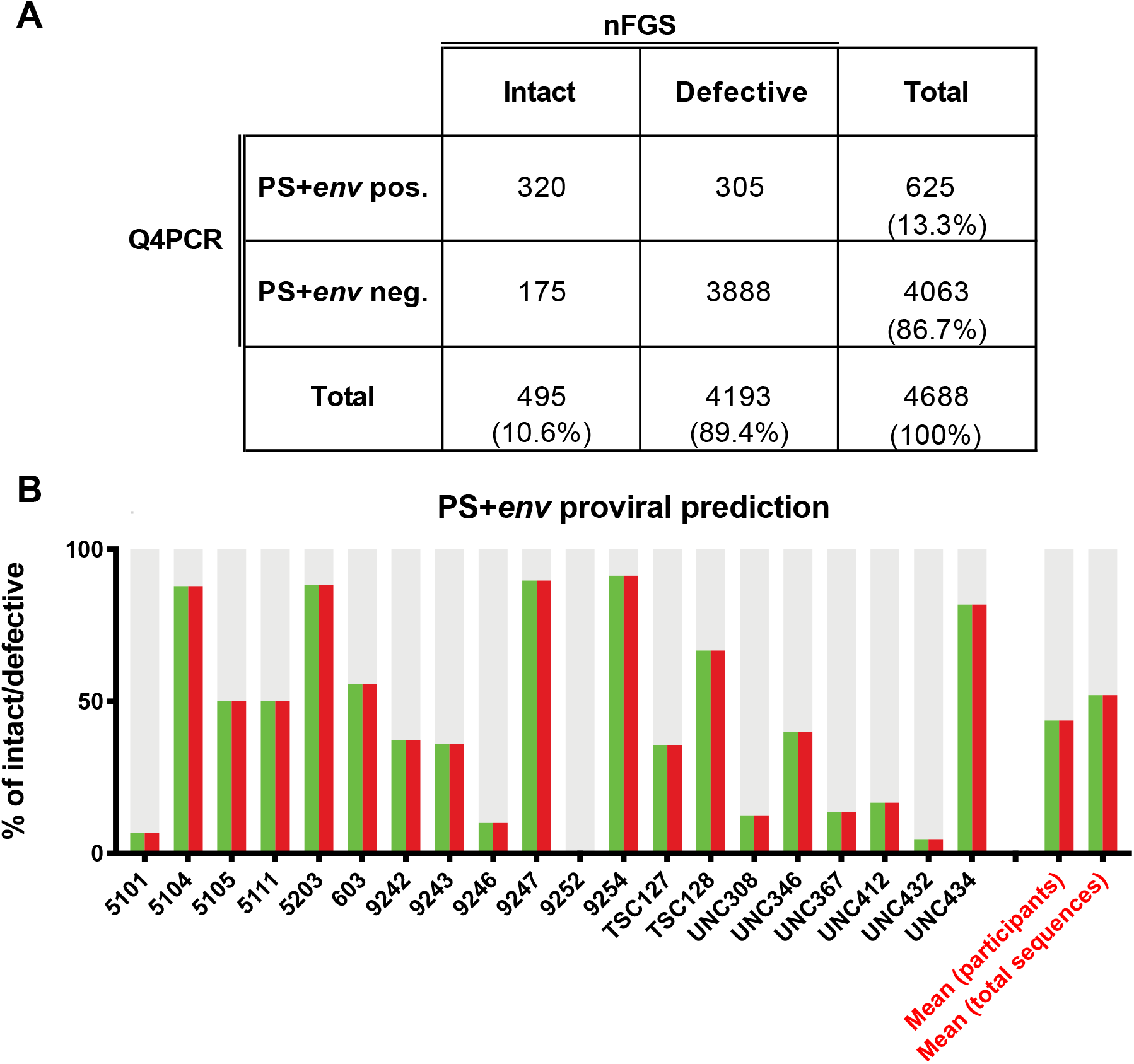
Q4PCR PS and *env* intact proviral genome prediction. **(A)** 2-by-2 table shows the number of PS+*env* positive and PS+*env* negative samples by Q4PCR and the corresponding classification as intact or defective proviral genomes based on near full-length sequencing results. To improve signal-to-background we excluded 29 sequences with Ct values greater than 38 for either the PS or *env* primer/probe set. **(B)** PS+*env* proviral sequence prediction. Q4PCR sequencing data were used to assess the fraction of true intact sequences out of all samples that were amplified by the Q4PCR PS and *env* primer/probe set combination on an individual participant basis. The graph shows the percentage of proviruses detected by the Q4PCR PS+env primer/probe combination that were truly intact by nFGS for 20 individuals with at least 5 PS+*env*-positive sequences. The predictive value for intact proviruses is colored in green (PS) and red (*env*). The defective fraction is shown in gray. The right two bar graphs depict the mean percentage of PS+*env*-positive samples that were truly intact by nFGS in a pooled analysis of all sequences (first bar graph to the right) or averaged across all 20 participants (second bar graph to the right), respectively.

Of all 625 PS+*env*-positive samples 320 or only 51.2% were actually identified as intact proviral genomes on a sequence level. To assess the percentage of PS+*env*-positive samples that were truly intact by nFGS on an individual participant basis, we analyzed individuals with at least 5 PS+*env*-positive sequences (Table S4). Importantly, and in line with our previous study the percent of PS+*env*-positive proviruses that were truly intact by nFGS appeared to vary between individuals (Fig. 4B) (9). PS+*env*+ proviruses are scored as intact by IPDA. However, we found that the frequencies of PS+*env*-positive proviruses detected by Q4PCR were a median of 6-fold (interquartile range 2, 17) lower than IPDA measurements (Table S2). Further, after sequence verification, Q4PCR values for intact viruses were substantially lower (by a median of 19-fold) than IPDA. Both of these results suggest that limitations of long-distance PCR amplification in Q4PCR as well as the IPDA misidentification of some PS and *env* amplifiable sequences as intact contribute to differences in the quantification of the intact proviral reservoir.

## Discussion

We compared the results of some of the available HIV-1 reservoir assays including Q4PCR and IPDA, because quantitation and characterization of the HIV reservoir informs efforts to develop an HIV cure (1). The great majority of integrated HIV proviral DNA is defective. Outgrowth assays quantitate the replication-competent reservoir but they are labor intensive and not scalable to large clinical studies. In addition, they do not capture all replication-competent viruses. The IPDA and Q4PCR were designed to address some of these issues. The IPDA utilizes ddPCR to measure proviruses that are PS+*env*-positive, which together exclude the great majority of defective proviruses with defects detectable by nFGS. This assay enables much more specific quantification of intact proviral DNA than single-probe assays. However, the PS and *env* probe combination is not entirely predictive of a fully intact provirus (8, 9). Q4PCR improves the sensitivity and specificity of PCR based assays for intact HIV DNA using a combination of qPCR and sequencing (9). The Q4PCR assay employs four different qPCR probes for PS, *env*, *pol*, and *gag* in a multiplex reaction to detect the HIV-1 genome in limiting dilution samples. Samples positive for 2 or more probes are subjected to near-full-length sequencing to verify the intactness of the provirus, enabling a very high specificity for intact proviruses. However, this assay is not scalable for large number of samples precisely because it requires limiting dilution PCR and full-length proviral DNA sequencing.

Our results show that Q4PCR and IPDA are positively correlated but differ in that IPDA identifies a higher number of intact proviruses than Q4PCR. Several possibilities, that are not mutually exclusive, may explain why intact proviral frequency estimates from Q4PCR were a median of 19-fold lower than those from IPDA (9).

First, limitations of long-distance PCR amplification in Q4PCR may have a positive bias toward amplification and subsequent quantification of defective proviruses due to their shorter length from large deletions (15). Inefficient amplification may account for the high degree of variation between the number of estimated intact genomes measured by different nFGS assays (4, 5, 9, 16–20) and it may explain why intact proviruses are not detected by Q4PCR in approximately 20% of infected individuals. In addition, even after correction for defects missed by the IPDA amplicons, intact provirus values measured by IPDA are substantially higher that those obtained by Q4PCR (Table S2).

A second factor that may contribute to higher intact proviral measurements in the IPDA versus Q4PCR is cell normalization methods. The IPDA normalizes proviral frequencies to cell input by quantifying host genome input using a parallel ddPCR measurement of the diploid host gene RPP30, while Q4PCR normalizes intact proviral numbers to the cell input based on DNA equivalents after CD4 isolation and DNA extraction, as measured by DNA fluorometry (Qubit).

The third independent factor that may contribute to higher intact proviral estimates from the IPDA may be misidentification of some amplifiable sequences as intact by the PS and *env* probe combination. Sequencing results from Q4PCR provides some insights into assay performance of the PS+*env* primer/probe combination. In this large sequencing study (506 intact and 4211 defective proviral sequences recovered), we found that the Q4PCR PS and *env* probe combination excluded 93% of defective proviruses in this dataset, consistent with the original report (8). With regard to the prediction of intact proviruses by the PS and *env* primer/probe combination, we observed that approximately 51% of the proviruses identified by the Q4PCR PS and *env* primer/probe combination are truly intact by nFGS. This is somewhat less than the 70% estimate from Bruner and colleagues based on analysis of 431 near full genome HIV-1 sequences (2.4% or 10 of which were intact) obtained by single genome analysis from 28 HIV-1-infected adults (8). As observed for Q4PCR, it is also conceivable that the precision of the IPDA PS and *env* probes to identify intact proviruses varies across individuals and should be considered in the interpretation of IPDA results. However, sequencing of proviruses in IPDA PS+*env* double positive droplets is needed to confirm this potential limitation (21), as our assessment of the frequency of PS+*env*-positive proviruses that are truly intact may be limited by the efficiency of nFGS. Despite a variable ability to predict intact proviral genomes, the IPDA PS and *env* probe combination appears to measure the decay of intact proviruses, as evidenced by the differential decay of intact versus defective proviruses (15, 22). Therefore, the IPDA is a useful high-throughput assay to establish an upper limit on the frequency of provirus that may be replication-competent.

Taken together, the IPDA affords a higher throughput capacity, but the combination of PS and *env* probes can both misclassify defective sequences and fail to detect intact ones due to sequence polymorphisms. In contrast, the sequencing information in Q4PCR enables a very high specificity for intact proviruses but it is not a high throughput assay, and Q4PCR quantitation of sequence-confirmed intact proviruses may be limited by the inefficiency of nFGS. Both assays are subject to probe amplification failures and HIV-1 subtype variation (11, 23), though Q4PCR may be less sensitive to this because of the use of 4 probes and subsequent sequencing of positive wells. Notably, both assays require minimal cell input relative to classical QVOA which is a clear advantage for clinical trials.

In conclusion, the differences across reservoir assays highlight that the predominantly defective and highly polymorphic proviral landscape makes the measurement of virus that is likely to cause rebound upon ART-cessation extremely challenging. Nonetheless, the IPDA and Q4PCR both represent major advances in accurately quantifying and characterizing the replication competent HIV reservoir. When used together, the two assays provide a deeper understanding of the reservoir than either assay alone and bracket true reservoir size.

## Materials and Methods

### CD4 T Cell Isolation

Cryopreserved PBMCs were viably thawed and magnetically negatively selected for CD4 T cells using either StemCell (cat#17952) or Miltenyi Biotech (cat# 130-096-533) kits.

### Intact proviral DNA assay

DNA was extracted from total CD4 T cells using the Qiagen QiaAmp DNA extraction kit as described, including the RNAse A step (8). DNA was measured on the NanoDrop 1000 Spectrophotometer (Thermo Fisher Scientific). 12-16 replicate wells of up to 800 ng DNA each were combined with 2X ddPCR Supermix for Probes (no dUTP, Bio-Rad) and primer/probe sets for regions in PS and *env* that favors amplification of intact proviral genomes, as described (3). Final concentrations of 750 nM of each primer and 250 nM of each probe (FAM/MGB for packaging signal probe; VIC/MGB for intact *env* probe and unlabeled/MGB for hypermutated exclusionary *env* probe) were used (Thermo Fisher Scientific). Droplets were generated on the Automated Droplet Generator (Bio-Rad), and subsequently underwent thermocycling on the C1000 Touch™ Thermal Cycler with a 96–Deep Well Reaction Module (Bio-Rad Cat#1851197). Thermal cycling conditions with a 2°C ramp rate were: 10 minutes at 95°C followed by 45 cycles (30 seconds at 94°C and 60 seconds at 59°C per cycle) and 10 minutes at 98°C, followed by incubation at 12°C as described (6). Droplets were read on the QX200 Droplet Reader (Bio-Rad Cat#1864003) using QuantaSoft Software Version 1.7.4.0917, with the exception of 2 samples, one (5114) with a confirmed PS polymorphism (Table S2), that made gating difficult that were read on the QX100 Droplet Reader. Replicate wells were merged and analyzed in the QuantaSoft Analysis Pro Software.

As described by Bruner and colleagues, a correction for DNA shearing was applied based on a separate, duplicate measurement of two regions of the diploid host gene RPP30 (4). RPP30 amplicons were spaced to match the distance between the PS and *env* IPDA HIV amplicons in order to quantify and correct for DNA shearing that occurs between the PS and *env* amplicons during the DNA extraction. Primer and probe sequences are available in Table S5. DNA shearing indices were highly consistent with other reports of IPDA measurements (Median 0.35, IQR 0.33-0.37) (Table S2) (8, 11). The RPP30 reaction was also used to normalize for cell input as described: for each individual, a median of 1.01×10^6^ (IQR 8.35 ×10^5^ – 1.16 ×10^6^) CD4 T cell genome equivalents (2 copies of the RPP30 gene) were assayed (8).

Buffer only no template controls (NTC) as well as DNA from HIV seronegative donors were used as a guide to set thresholds for positive droplets on each digital PCR plate. In our hands, we observed occasional low-amplitude false positive droplets in NTC wells at a rate less than or equal to 5 copies/million CD4 T cells, depending on the run. Based on the observed false positive rate and the subsampling constraints of digital PCR, IPDA proviral frequencies were censored at 5 copies/million CD4 T cells. In some individuals, a proviral polymorphism in primer/probe binding regions resulted in a droplet spread phenotype that was difficult to set thresholds for because of insufficient droplet separation from the negative population (Fig. S3, Table S2). These 4 individuals were included in subsequent analyses (Table S2). In some individuals, a proviral polymorphism precluded PCR amplification of one of the IPDA amplicons (PS or *env*) (Table S2). These individuals’ IPDA data were excluded from the analyses.

### gag HIV DNA assay

DNA was extracted from total CD4 T cells using the Qiagen QiaAmp DNA extraction kit. Up to 8 replicate wells of up to 500 ng of DNA was added to ddPCR™ Supermix for Probes (No dUTP) (Bio-Rad) with a primer/probe set overlapping the LTR-*gag* junction designed to measure group-M HIV-1 proviral load (10). 900 nM of each primer and 250 nM of probe was used. Droplets were generated on the Automated Droplet Generator (Bio-Rad), and subsequently underwent thermocycling with a 2°C ramp rate for 10 minutes at 95°C for 45 cycles (30 seconds at 94°C and 60 seconds at 59°C per cycle) and 10 minutes at 98°C prior to reading on the QX200 droplet reader. RPP30 reactions were used to normalize for cell input. In 2 individuals, no positive droplets were identified. These were considered PCR amplification failures and excluded from further analysis because positive droplets were identified using the partially overlapping IPDA PS primer/probe set in the same 2 individuals.

### Q4PCR

Q4PCR was performed as previously described (9). Briefly, genomic DNA from 1-5 million total CD4+ T cells was isolated using the Gentra Puregene Cell Kit (Qiagen) or phenol-chloroform and the DNA concentration was measured by Qubit High Sensitivity Kit (Thermo Fisher Scientific). Next, an outer PCR reaction (NFL1) was performed on genomic DNA at a single-copy dilution (previously determined by gag limiting dilution) using outer PCR primers BLOuterF (59-AAATCTCTA GCAGTGGCGCCCGAACAG-39) and BLOuterR (59-TGAGGGATCTCT AGTTACCAGAGTC-39, (5)). Undiluted 1-μl aliquots of the NFL1 PCR product were subjected to a Q4PCR reaction using a combination of four primer/probe sets that target conserved regions in the HIV-1 genome. Each primer/probe set consists of a forward and reverse primer pair as well as a fluorescently labeled internal hydrolysis probe as previously described (9): PS forward, 59-TCTCTCGACGCA GGACTC-39; reverse, 59-TCTAGCCTCCGCTAGTCAAA-39; probe, 59-/Cy5/TTTGGCGTA/TAO/CTCACCAGTCGCC-39/IAbRQSp (Integrated DNA Technologies); *env* forward, 59-AGTGGTGCAGAGAGAAAAAAGAGC-39; reverse, 59-GTCTGGCCTGTACCGTCAGC-39; probe, 59-/VIC/CCTTGGGTTC-TTGGGA-39/MGB (Thermo Fisher Scientific); gag forward, 59-ATGTTTTCAGCATTATCAGAAGGA-39; reverse, 59-TGCTTGATGTCCCCCCACT-39; probe, 59-/6-FAM/CCACCCC-AC/ZEN/AAGATTTAAACACCATGCTAA-39/IABkFQ (Integrated DNA Technologies); and pol forward, 59-GCA CTTTAAATTTTCCCATTAGTCCTA-39; reverse, 59-CAAATTTCTACT AATGCTTTTATTTTTTC-39; probe, 59-/NED/AAGCCAGGAATGGA-TGGCC-39/MGB (Thermo Fisher Scientific). Each Q4PCR reaction was performed in a 10-μl total reaction volume containing 5 μl TaqMan Universal PCR Master Mix. containing Rox (4304437; Applied Biosystems), 1 μl diluted genomic DNA, nuclease-free water, and the following primer and probe concentrations: PS, 675 nM of forward and reverse primers with 187.5 nM of PS internal probe; *env*, 90 nM of forward and reverse primers with 25 nM of *env* internal probe; gag, 337.5 nM of forward and reverse primers with 93.75 nM of gag internal probe; and pol, 675 nM of forward and reverse primers with 187.5 nM of pol internal probe. qPCR conditions were 94°C for 10 min, 40 cycles of 94°C for 15 s, and 60°C for 60 s. All qPCR reactions were performed in a 384-well plate format using the Applied Biosystem QuantStudio 6 Flex Real-Time PCR System. qPCR data analysis was performed as previously described (9). Generally, samples showing reactivity with two or more of the four qPCR probes were selected for a nested PCR reaction (NFL2). The NFL2 reaction was performed on undiluted 1-μl aliquots of the NFL1 PCR product. Reactions were performed in a 20-μl reaction volume using Platinum Taq High Fidelity polymerase (Thermo Fisher Scientific) and PCR primers 275F (59-ACAGGGACCTGAAAGCGAAAG-39) and 280R (59-CTAGTT ACCAGAGTCACACAACAGACG-39, (5)) at a concentration of 800 nM. Library preparation and sequencing was performed as previously described (9).

As previously described, HIV-1 sequence assembly was performed by our in-house pipeline (Defective and Intact HIV Genome Assembler), which is capable of reconstructing thousands of HIV genomes within hours via the assembly of raw sequencing reads into annotated HIV genomes (24). The steps executed by the pipeline are described briefly as follows. First, we removed PCR amplification and performed error correction using clumpify.sh from BBtools package v38.72 (https://sourceforge.net/projects/bbmap/). A quality control check was performed with Trim Galore package v0.6.4 (https://github.com/FelixKrueger/TrimGalore) to trim Illumina adapters and low-quality bases. We also used bbduk.sh from BBtools package to remove possible contaminant reads using HIV genome sequences, obtained from Los Alamos HIV database, as a positive control. We used a k-mer–based assembler, SPAdes v3.13.1, to reconstruct the HIV-1 sequences. The longest assembled contig was aligned via BLAST to a database of HIV genome sequences, obtained from Los Alamos, to set the correct orientation. Finally, the HIV genome sequence was annotated by aligning against Hxb2 using BLAST. Sequences with double peaks, i.e., regions indicating the presence of two or more viruses in the sample (cutoff consensus identity for any residue <70%), or samples with a limited number of reads (empty wells ≤500 sequencing reads) were omitted from downstream analyses. In the end, sequences were classified as intact or defective(24).

### Quantitative Viral outgrowth assays

The quantitative and qualitative viral outgrowth assay (Q2VOA) was performed as previously described (25, 26). In brief, isolated CD4+ T cells were activated with phytohemagglutinin and IL-2 (Peprotech) and cocultured with 1 × 10^6^ irradiated PBMCs from a healthy donor in 24-well plates. After 24 h, PHA was removed and 0.1 × 10^6^ MOLT-4/CCR5 cells were added to each well. Cultures were maintained for two weeks, splitting the cells seven days after the initiation of the culture and every other day after that. Positive wells were detected by measuring p24 by ELISA. The frequency of latently infected cells was calculated through the IUPM algorithm developed by the Siliciano laboratory (http://silicianolab.johnshopkins.edu) as previously described (25).

The quantitative viral outgrowth assay (QVOA) was performed on resting CD4 T cells for participants from the UNC cohort. Lymphocytes were obtained by continuous-flow leukapheresis and isolation of resting CD4^+^ T cells. Recovery and quantification of replication-competent virus was performed as described elsewhere (27). In general, approximately 50 million resting CD4^+^ T cells were plated in replicate limiting dilutions of 2.5 million (18 cultures), 0.5 million (6 cultures), or 0.1 million (6 cultures) cells per well, activated with phytohemagglutinin (Remel), a 5-fold excess of allogeneic irradiated PBMCs from a seronegative donor, and 60 U/mL interleukin 2 for 24 hours. Cultures were washed and cocultivated with CD8-depleted PBMCs collected from selected HIV-seronegative donors screened for adequate CCR5 expression. Culture supernatants were harvested and assayed for virus production by p24 antigen-capture enzyme-linked immunosorbent assay (ABL). Cultures were scored positive if p24 was detected at day 15 and was increased in concentration at day 19. The number of resting CD4^+^ T cells in infected units per billion was estimated using a maximum likelihood method (28).

## Data availability

Proviral sequences have been deposited in GenBank with the accession nos. MN090188-MN090943, MT189273-MT191207 and MW059111-MW063110.

## Acknowledgments

We thank all study participants who devoted time to our research; the Rockefeller University Hospital Research support office and nursing staff; all members of the M.C.N. laboratory for discussions and M. Jankovic for laboratory support; C.G. was supported by the Robert S. Wennett Post-Doctoral Fellowship, in part by the National Center for Advancing Translational Sciences (National Institutes of Health Clinical and Translational Science Award program, grant UL1 TR001866) and by the Shapiro-Silverberg Fund for the Advancement of Translational Research. E.S. was supported by the Swiss National Science Foundation (Grant number P1ZHP3_188135). This work was supported by the Bill and Melinda Gates Foundation (Collaboration for AIDS Vaccine discovery grants OPP1092074) and the National Institutes of Health (grants UM1 AI100663 and R01AI129795 to M.C. Nussenzweig, 1U01AI129825 to M. Caskey, UM1AI126619 to D.M. Margolis, R01AI134363 to N.M. Archin, P30AI050410, and F30AI145588 to S.D. Falcinelli); the Einstein–Rockefeller–CUNY Center for AIDS Research (grant 1P30AI124414-01A1); BEAT-HIV Delaney (grant UM1 AI126620 to M. Caskey); and the Robertson Fund. M.C. Nussenzweig is a Howard Hughes Medical Institute Investigator.

## Author contributions

CG, SDF, JCCL, MC, NMA, DMM, and MCN designed experiments. CG, SDF, ES, JR, RM, JK, KSJ, BA performed experiments. CG, SDF, ES, JCCL, TYO, VR MC, NMA, DMM, and MCN analyzed the data. CB and JDK collected clinical data and recruited participants. CG, SDF, NMA, RFS, DMM, and MCN wrote the manuscript.

---

**Fig. S1:**
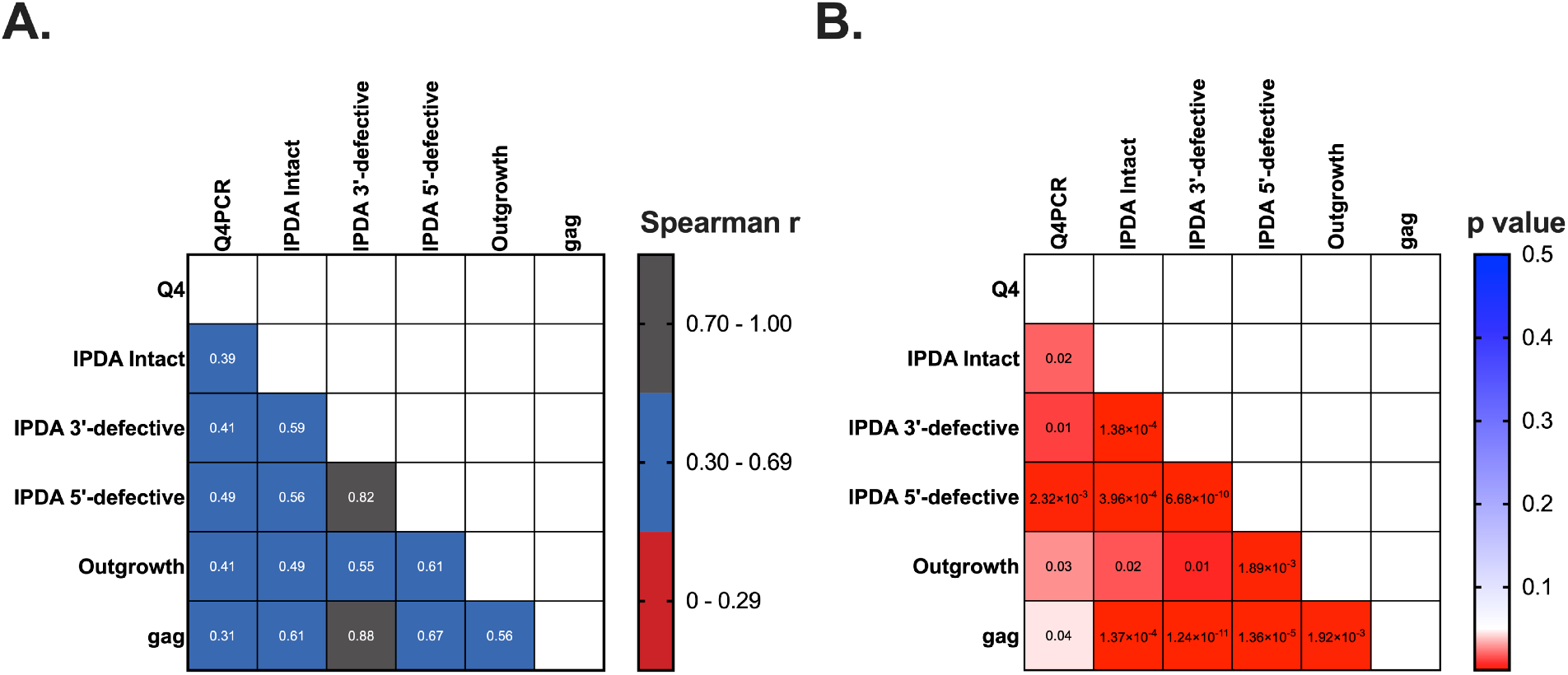
Spearman correlation and p value matrices comparing all available reservoir measurements. p values are unadjusted. Left censored measurements were included: for Q4PCR, when no intact proviruses were recovered a value of 0 was used and for IPDA, a left-censor of 5 copies/million CD4 T cells was used as described in the Methods. Data points were excluded when an IPDA amplification failure was present.

**Fig. S2:**
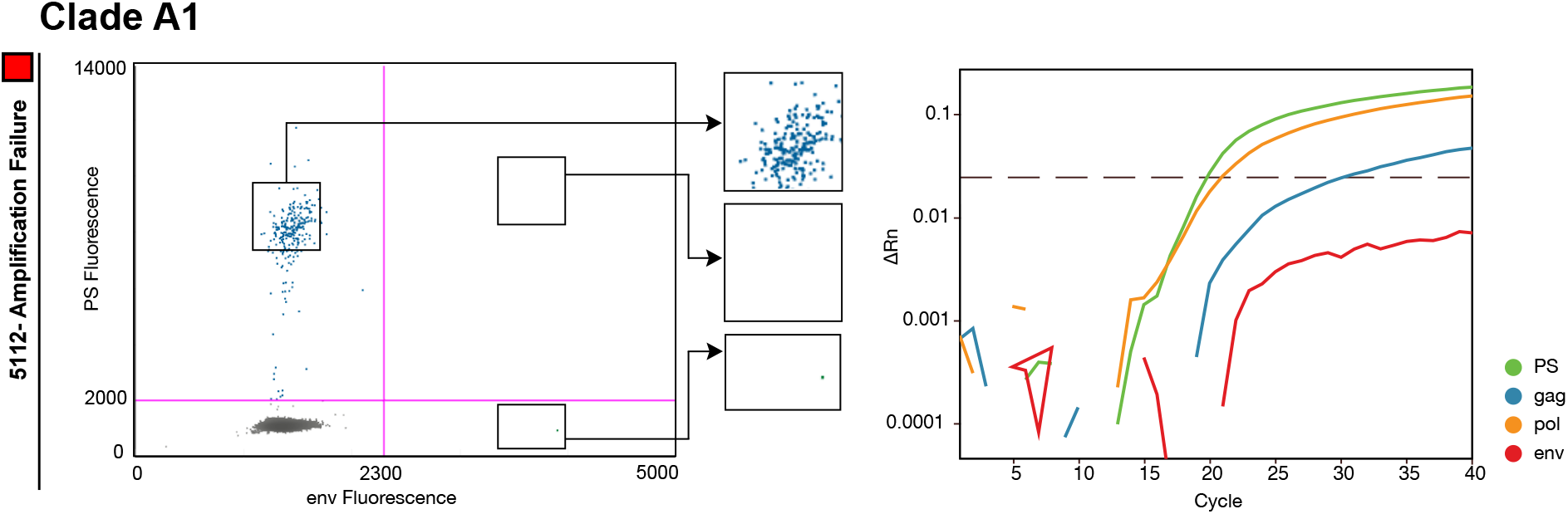
Non-clade B signal quality. Example of an amplification failure of the *env* signal in IPDA for participant 5112. The corresponding Q4PCR signal of an intact proviral genome also shows an *env* signal failure but at the same time a clear signal for the PS and *pol* probe demonstrating less susceptibility to non-clade B viral genotypes for Q4PCR.

**Fig. S3:**
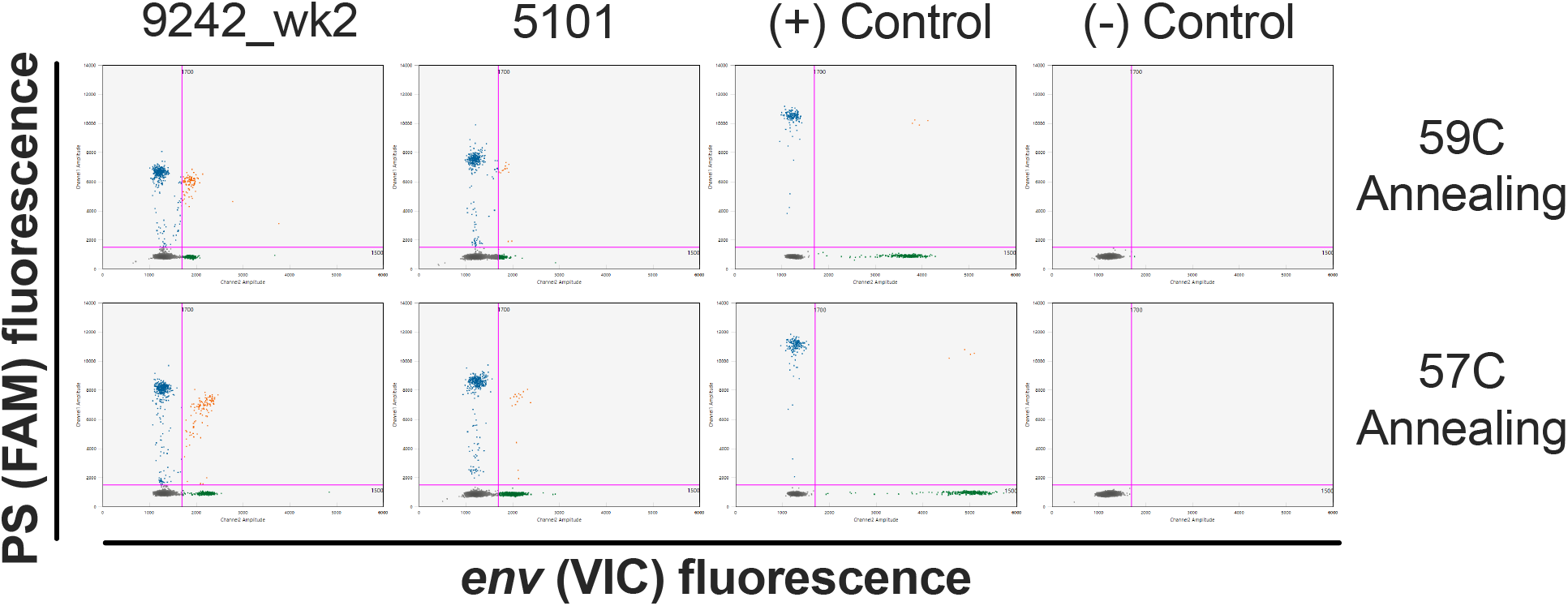
Improved droplet separation for the IPDA with 57°C versus 59°C annealing temperature. Droplet amplitude improved, especially for the *env*/VIC amplicon, with a PCR annealing temperature of 57°C versus 59°C without compromising the ability of the *env* hypermutated probe to discriminate hypermutated sequences (data not shown).

**Table S1:** Participant Characteristics. Individual participant demographics and baseline clinical characteristics Age, sex, race/ethnicity, time since HIV diagnosis, time since ART initiation, ART regimen at time of donation, and estimated CD4 nadir are provided for the n=39 ART-suppressed participants studied.

| ID | Age | Sex | Race | Years since |  | ART at Screening | Estimated CD4 nadir |
| --- | --- | --- | --- | --- | --- | --- | --- |
|  |  |  |  | HIV-1 dx | first ART |  |  |
| 9241 | 40 | M | White/Hisp | 6 | 7 | EVG/cobi/TDF/FTC | 500 |
| 9242 | 43 | M | White/Hisp | 3 | 3 | EVG/cobi/ TDF/FTC | 450 |
| 9243 | 29 | M | Amer Indian/Hisp | 5 | 5 | RPV/TDF/FTC | 350 |
| 9244 | 36 | M | Amer Indian/Non-Hisp | 9 | 5 | EFV/TDF/FTC | 730 |
| 9246 | 30 | M | Black | 5 | 5 | EVG/cobi/ TAF/FTC | 500 |
| 9247 | 31 | M | Black | 6 | 6 | EVG/cobi/ TAF/FTC | 600 |
| 9252 | 51 | F | Black | 11 | 11 | EFV/TDF/FTC | 270 |
| 9254 | 48 | M | White | 21 | 21 | EVG/cobi/ TAF/FTC | 590 |
| 9255 | 30 | M | White | 5 | 4 | EVG/cobi/ TAF/FTC | 779 |
| B207 | 48 | M | White/Hisp | 12 | 11 | EVG/TDF/FTC | not available |
| 603 | 45 | M | White/Hisp | 14 | 12 | EFV/TDF/FTC | 300 |
| 605 | 38 | M | White/Hisp | 17 | 16 | RPV/TDF/FTC | 372 |
| 5104 | 35 | M | Black | 7 | 7 | BIC/TAF/FTC | 400 |
| 5106 | 31 | M | Black | 6 | 6 | EVG/COBI/FTC/TDF | >350 |
| 5108* | 52 | M | White | 10 | 10 | EVG/COBI/FTC/TAF | >300 |
| 5112* | 30 | M | White | 6 | 5 | EVG/COBI/FTC/TAF | 350 |
| 5101 | 52 | M | Multiple/Hisp | 12 | 12 | TDF/FTC/EFV | not available |
| 5114 | 54 | M | Black | 15 | 15 | ABC/DTG/3TC | 300 |
| 5105 | 49 | F | Black | 29 | 29 | FTC/RPV/TDF | 200 |
| 5203 | 59 | M | White | 23 | 21 | DTG/RPV | 500 |
| 5111 | 55 | M | White | 20 | 16 | EVG/COBI/FTC/TAF | 400 |
| 5115 | 36 | M | Black | 6 | 6 | FTC/TAF/DTG | not available |
| TSC-124 | 52 | M | Black | 10 | 10 | EFV/TDF/FTC | not available |
| TSC-127 | 57 | F | Multiple/Non-Hisp | 20 | 20 | EVG/COBI/FTC/TAF | not available |
| TSC-125 | 43 | M | White | 18 | 11 | EVG/COBI/FTC/TDF | not available |
| TSC-131 | 45 | M | Multiple | 1 | 1 | EFV/TDF/FTC | not available |
| TSC-128 | 43 | M | N/A / Non-Hisp | 18 | 18 | FTC/RPV/TDF | not available |
| UNC-432 | 51 | M | White | 11 | 11 | ABC/DTG/3TC | 403 |
| UNC-308 | 53 | M | White | 22 | 16 | DRV/ABC/3TC/NEV/RTV | 81 |
| UNC-346 | 51 | M | White | 10 | 9 | EVG/cobi/FTC/TAF | 168 |
| UNC-367 | 28 | M | White | 6 | 5 | DRV/cobi/FTC/TAF | 168 |
| UNC-412 | 31 | M | White | 8 | 8 | ABC/DTG/3TC | 354 |
| UNC-434 | 45 | M | Other/Hisp | 6 | 6 | BIC/FTC/TAF | 526 |
| UNC-404 | 25 | M | Black | 2 | 2 | ABC/DTG/3TC | 308 |
| UNC-336 | 53 | M | Black | 22 | 20 | EVG/cobi/FTC/TAF | 462 |
| UNC-406,425** | 43/44 | M | Black | 1,2 | 1,2 | ABC/DTG/3TC | 713 |
| UNC-397 | 34 | M | Black | 10 | 4 | EVG/cobi/FTC/TAF | not available |
| UNC-437 | 42 | F | Black | 9 | 8 | EVG/cobi/FTC/TAF | 350 |
| UNC-458* | 42 | F | Black | 11 | 11 | BIC/FTC/TAF | not available |

EVG - elvitegravir, cobi - cobicistat, TDF - tenofovir disoproxil fumarate, FTC - emtricitabine, RTV - ritonavir, ABC - abacavir, 3TC - lamivudine, DTG - dolutegravir. BIC - bictegravir RPV - rilpivirine, EFV - efavirenz, TAF - tenofovir alafenamide fumarate, DRV - darunavir, NEV- nevirapine \*These participants harbored non-clade B viruses. \*\*This individual provided 2 donations at 1 year and 2 years following ART initiation.

**Table S2:**
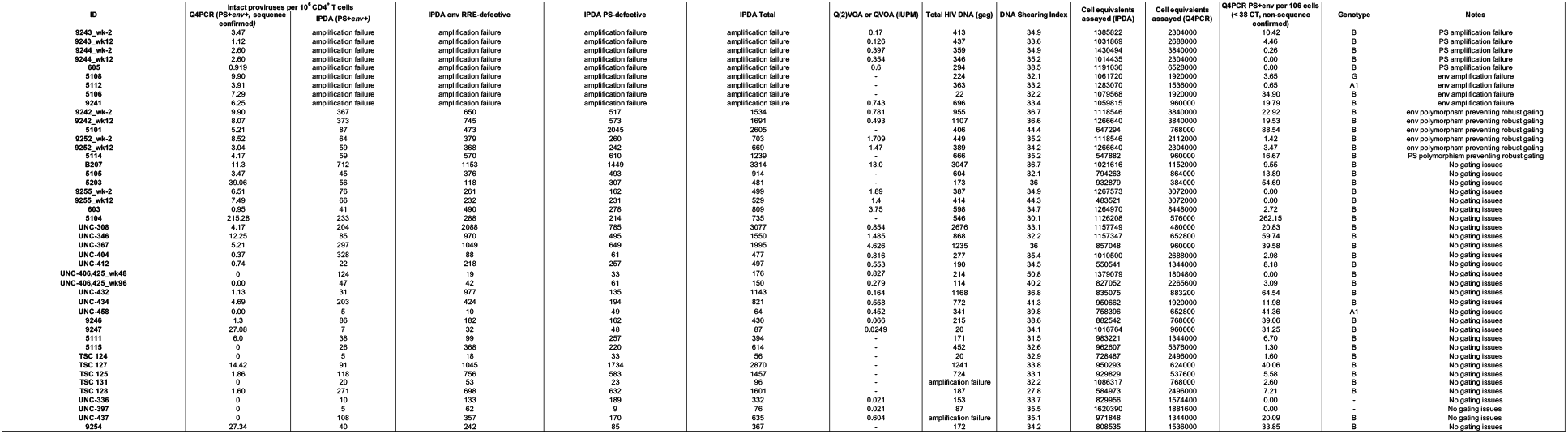
Quantitative Analysis. Table S2 shows the results of the quantitative reservoirs measurements for the 39 ART-suppressed participants studied.

**Table S3:** Sequences per participant. Table S3 lists the number of sequences obtained by Q4PCR for each of the 39 participants

| ID | sequenced | intacts | defective |
| --- | --- | --- | --- |
| 5101 | 156 | 4 | 152 |
| 5104 | 158 | 124 | 34 |
| 5105 | 120 | 3 | 117 |
| 5106 | 20 | 14 | 6 |
| 5108 | 76 | 19 | 57 |
| 5111 | 101 | 8 | 93 |
| 5112 | 74 | 6 | 68 |
| 5114 | 60 | 4 | 56 |
| 5115 | 17 | 0 | 17 |
| 5203 | 29 | 15 | 14 |
| 603 | 461 | 8 | 453 |
| 605 | 90 | 6 | 84 |
| 9242_wk-2 | 318 | 38 | 280 |
| 9242_wk12 | 303 | 31 | 272 |
| 9243_wk-2 | 195 | 8 | 187 |
| 9243_wk12 | 93 | 3 | 90 |
| 9244_wk-2 | 155 | 10 | 145 |
| 9244_wk12 | 127 | 6 | 121 |
| 9246 | 32 | 1 | 31 |
| 9247 | 49 | 26 | 23 |
| 9241 | 134 | 6 | 128 |
| 9252_wk-2 | 213 | 19 | 194 |
| 9252_wk12 | 241 | 7 | 234 |
| 9254 | 98 | 42 | 56 |
| 9255_wk-2 | 306 | 20 | 286 |
| 9255_wk12 | 261 | 24 | 237 |
| B207 | 57 | 13 | 44 |
| TSC124 | 7 | 0 | 7 |
| TSC125 | 20 | 1 | 19 |
| TSC127 | 136 | 9 | 127 |
| TSC128 | 41 | 4 | 37 |
| TSC131 | 1 | 0 | 1 |
| UNC308 | 122 | 2 | 120 |
| UNC336 | 0 | 0 | 0 |
| UNC346 | 155 | 8 | 147 |
| UNC367 | 143 | 5 | 138 |
| UNC397 | 0 | 0 | 0 |
| UNC404 | 35 | 1 | 34 |
| UNC-406,425_wk48 | 0 | 0 | 0 |
| UNC-406,425_wk96 | 5 | 0 | 5 |
| UNC412 | 15 | 1 | 14 |
| UNC432 | 43 | 1 | 42 |
| UNC434 | 46 | 9 | 37 |
| UNC437 | 3 | 0 | 3 |
| UNC458 | 1 | 0 | 1 |

**Table S4:** PS+env sequence prediction. Tables S4 shows the number and classification (intact vs. defective) of PS+*env* positive samples for all participants with at least 5 PS+*env* positive samples.

| IDs | total | Intact | def | % intact |
| --- | --- | --- | --- | --- |
| 5101 | 58 | 4 | 54 | 6.9% |
| 5104 | 140 | 123 | 17 | 87.9% |
| 5105 | 6 | 3 | 3 | 50.0% |
| 5111 | 8 | 4 | 4 | 50.0% |
| 5203 | 17 | 15 | 2 | 88.2% |
| 603 | 9 | 5 | 4 | 55.6% |
| 9242 | 137 | 51 | 86 | 37.2% |
| 9243 | 25 | 9 | 16 | 36.0% |
| 9246 | 10 | 1 | 9 | 10.0% |
| 9247 | 29 | 26 | 3 | 89.7% |
| 9252 | 11 | 0 | 11 | 0.0% |
| 9254 | 46 | 42 | 4 | 91.3% |
| TSC127 | 14 | 5 | 9 | 35.7% |
| TSC128 | 6 | 4 | 2 | 66.7% |
| UNC308 | 8 | 1 | 7 | 12.5% |
| UNC346 | 20 | 8 | 12 | 40.0% |
| UNC367 | 22 | 3 | 19 | 13.6% |
| UNC412 | 6 | 1 | 5 | 16.7% |
| UNC432 | 22 | 1 | 21 | 4.5% |
| UNC434 | 11 | 9 | 2 | 81.8% |

**Table S5:**
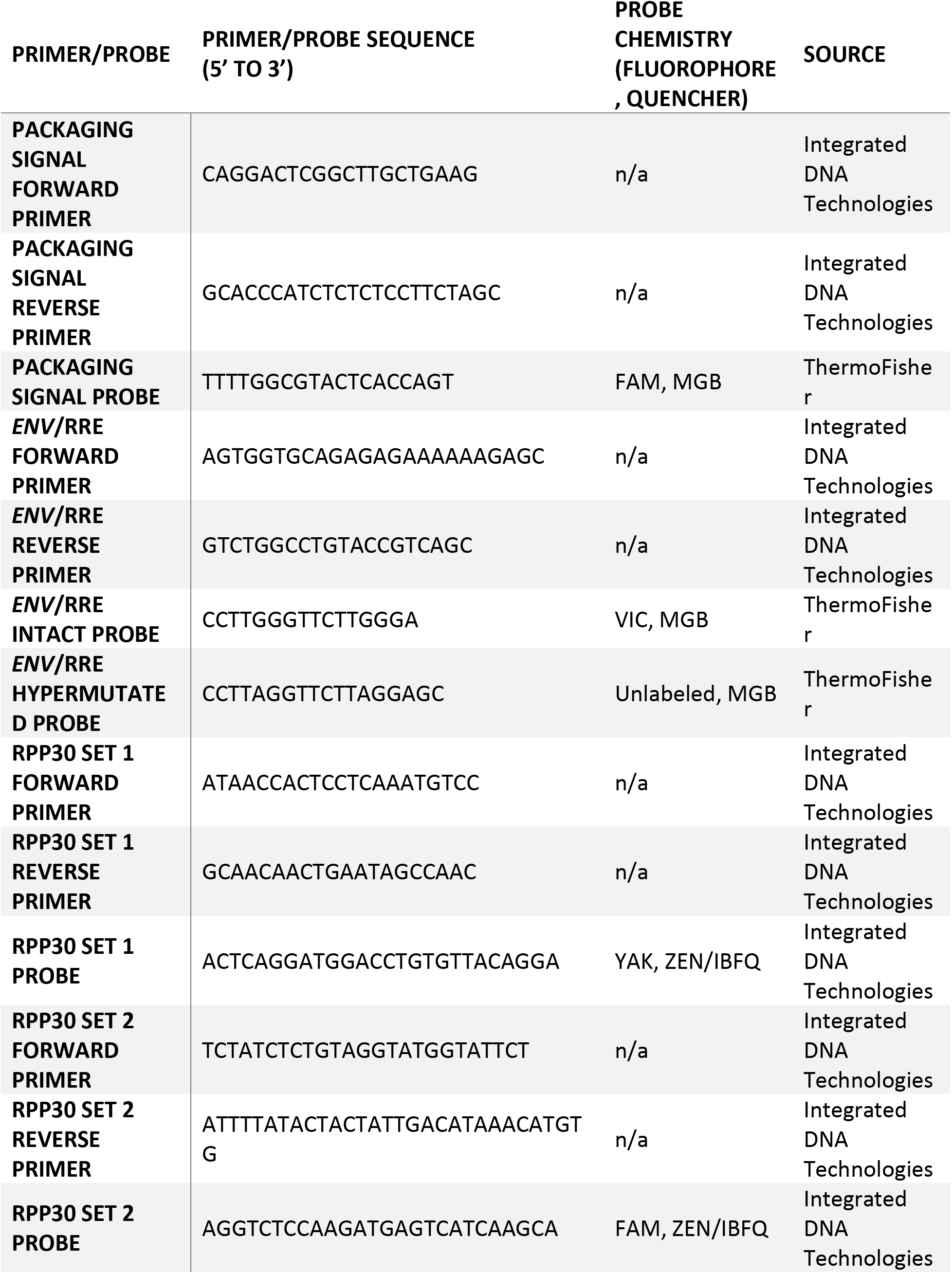
Primer and probe sequences for the Intact Proviral DNA Assay. Detailed parameters for the Intact Proviral DNA Assay used in the current study. This assay was originally designed by Bruner and colleagues (8).

